# A switch from histone methyltransferase-EZH2 to demethylase KDM6A activity marks reinitiation of proliferation of drug treated colorectal cancer cells

**DOI:** 10.1101/2025.11.04.686665

**Authors:** Subhashree Chatterjee, Ritika Jaiswal, Aniruddha Roy, Shibasish Chowdhury, Sudeshna Mukherjee, Rajdeep Chowdhury

## Abstract

Colorectal cancer (CRC) is one of the deadliest cancers, ranking third in cancer incidence worldwide. These tumor cells often adopt unique strategies under drug stress to attain a reversible drug-tolerant state and evade cell death. However, the molecular adaptations associated with this transitory emergence of the drug-tolerant state remain elusive. Herein, epigenetic alterations often dictate such reversible dynamic changes, and this study aims to characterize the role of specific epigenetic modifiers governing CRC cell survival under drug pressure and their subsequent relapse. We observed that under drug stress there is a drastic increase in the histone-repressive mark-H3K27me3, linked to an enhanced expression of EZH2 driving transcriptional inhibition of cell proliferation-associated genes and a proliferative arrest. Interestingly, drug-induced oxidative stress increased the expression of the P65 protein, which was found to interact and regulate EZH2 expression. Quenching of ROS, drug vacation, or P65 inactivation compromised EZH2 activity concurrent with a re-initiation of cell proliferation. Interestingly, this reversal to proliferative state was associated with an elevated activity of the histone lysine demethylase-KDM6A. The promoter elements of the proliferative genes were now occupied by KDM6A instead of EZH2. Accordingly, a genetic knockdown or pharmacological inhibition of KDM6A *in vitro* not only resulted in increased cell death but also prevented emergence of the re-proliferative CRC cells. Furthermore, KDM6A inhibition in combination with chemotherapy drug, resulted in an increased tumor regression *in vivo*. Our study thus highlights the importance of KDM6A as a therapeutic target in preventing CRC growth and relapse which can have future therapeutic implications.

## 1. Introduction

Conventional chemotherapy is directed toward eliminating fast-proliferating tumor cells; however, existing reports suggest that a small population of tumor cells can survive this toxic drug stress. These persisters with time attain proliferative potential and can contribute towards subsequent relapse [1–3]. Given that recurrence after chemotherapy is one of the major obstacles to cancer therapy, identifying routes by which persisters can be obliterated is of key interest. Recent reports suggest that, unlike genetic changes, epigenetic alterations are dynamic, reversible in nature and are an appropriate adaptive strategy for the tumor cells surviving stress [4–7]. Therefore, dynamic regulation of chromatin architecture and associated transcriptional reprogramming hitherto absent before drug treatment is now acknowledged as a potent strategy acquired by the tumor cells to temporarily avert drug stress [8–10]. However, further studies are required to identify epigenetic factors that determine this transitory tolerant phase and relapse, and how it can be circumvented.

Among different epigenetic modifications at play under drug stress, histone modifications have recently gained the limelight [11,12]. The N-terminal tail of histones is prone to various post-translational modifications, which determine the transcriptional fate of many tumorigenesis-related genes as well as chromatin structures [13–15]. In this regard, histone H3 methylation, more specifically on lysine 27 (H3K27), is putatively one of the most abundant modifications that dictate the active or repressed state of a gene based on its mono, di or tri methylation status [16,17]. The chromatin accessibility through H3K27 methylation has been observed to be precisely monitored by the activity of two enzymes, the histone methyltransferase-EZH2 and histone lysine demethylase KDM6A, known as UTX [18–20]. Interestingly, both EZH2 and KDM6A mutation and their dysregulation are linked with various cancer types, acting as a repressor or an activator in a dual-faceted and/or context-dependent manner [21–23]. For example, EZH2 overexpression is frequently associated with tumor aggressiveness. In breast cancer, particularly in rapidly proliferating subtypes, such as triple-negative and ERBB2-overexpressing tumors, elevated EZH2 levels correlate with increased malignancy [24]. Notably, EZH2 overexpression has also been shown to promote neoplastic transformation of normal breast epithelial cells [25]. Beyond breast cancer, similar associations between EZH2 overexpression, tumor aggressiveness, and poor prognosis have been reported in various other cancers as well, including thyroid cancer, hepatocellular carcinoma, and oesophageal squamous cell carcinoma [26–28]. In contrast, recent reports also suggest loss of function mutation in EZH2, which in turn accelerates Kras-driven lung adenocarcinoma [29]. Furthermore, in diffuse midline gliomas, genetic ablation of EZH2 increased tumor growth through an inflammatory response. Interestingly, EZH2 gain-of-function mutation significantly reduced tumor incidence and increased tumor latency through oxidative phosphorylation/mitochondrial metabolic pathway [30]. Overall, EZH2 functions as a pro- or anti-tumorigenic molecule purely in a context-dependent manner, demanding more investigation into its functional modality. Herein, EZH2, the catalytic subunit of the polycomb repressive complex 2 (PRC2) family of proteins, canonically acts in association with its core subunits-SUZ12, EED and RBAP46/8 and facilitates the hierarchical recruitment of methyl group on the lysine 27 of histone H3 (H3K27me3) [31–33]. Also, the core PRC2 members associate with other proteins like AEBP2 and JARID2; these two proteins are known to synergistically enhance the catalytic activity of EZH2 [34].

While, in contrast to canonical EZH2 activity, the gene encoding KDM6A, located on the X chromosome, catalyses the removal of di or trimethylated H3K27 through its Jumonji C (JmjC) domain [35]. Interestingly, context-dependent regulation of KDM6A often raises concern in therapeutic implications. For example, an epigenetic switch triggered by KDM6A loss disrupts urothelial differentiation by disrupting FOXA1 target binding capability and promoting genome-wide redistribution of ATF3, leading to repression of FOXA1 targets, activation of cell-cycle genes, resulting in increased cellular proliferation [36]. Similarly, loss of KDM6A promotes squamous-like and metastatic pancreatic cancer in females [37]. Also, in breast epithelial cells, loss of KDM6A resulted in reduced responsiveness to anticancer therapies [38]. In contrast, existing reports also show that an upregulation in KDM6A is positively correlated with cellular proliferation as well as invasion in breast cancer patients, which is often linked to poor clinical outcome [39]. Furthermore, an elevated KDM6A expression is shown to promote the formation of glioblastoma stem cell populations by enhancing Notch signaling and sustaining a slow cell cycle, thereby contributing to tumor relapse [5]. Importantly, several upstream molecules or signaling pathways have been implicated in the regulation of both EZH2 or KDM6A. However, the tumor microenvironment potentially determines the molecules involved in their regulation, and this requires further exploration. In this regard, intracellular reactive oxygen species (ROS) and their downstream mediators deserve a special mention. Importantly, in a tumor microenvironment, redox imbalance is not only limited to DNA damage but also linked with abnormal intracellular signaling cascades leading to tumor heterogeneity and pathogenicity [40–42]. In this context, a stress-regulated transcription factor of the Rel family, NF-KB P65 plays a pivotal role in maintaining oxidative response and simultaneously regulating multiple aspects of oncogenesis, including proliferation and cell cycle [43,44]. Recent reports have indicated that NF-KB P65 also interacts with different histone modifiers under stress conditions, leading to chromatin remodelling and associated gene expression [45,46]. However, the role of such molecular regulators in dictating the action of the epigenetic factors like EZH2/KDM6A under drug stress is not fully understood.

Hence, although a myriad of reports highlight the regulation of key epigenetic modulators, it remains unclear how they are in turn regulated and how they temporally modulate gene expression in tumor cells, especially after drug stress. Here, in this study, we show that drug stress results in an increase in ROS and associated P65 signalling, resulting in enhanced nuclear translocation of EZH2, and induction of a repressed chromatin state with attenuated cell proliferation of colorectal cancer cells (CRC). However, drug withdrawal or ROS quenching results in a ‘switch on’ of KDM6A activity and abrogation of EZH2 function leading to re-proliferation of the tumor cells. Taken together, this study provides key insights into how epigenetic modulators like EZH2 and KDM6A function and can be involved in chemo tolerance and survival of cancer cells.

## 2. Materials and methods

### 2.1 Chemicals and reagents

Cisplatin (#232120), Trizol reagent (#T9424), Fetal bovine serum albumin (#F7524), RIPA Buffer (#R0278), Protease inhibitor cocktail (#P83409), Triton X 100 (#T8787), Tween 20 (#P1379-1L), Bradford reagent (#B6916), Acrylamide (#A9099) N’N Methylenebisacrylamide (#M7279), GSK J4 (#SML0701), Corning® Matrigel® Basement Membrane Matrix (#CLS354234) were obtained from Sigma Aldrich. Trypsin (#25300062), Antibiotic-Antimycotic (100X) (#15240062), and Opti-MEM (#31985070) were procured from Gibco. DPBS (#21300025), Lipofectamine 3000 (#L3000015), DNase 1, RNase free (#EN-0521), Annexin V, FITC-Conjugate (#A13199), MAGnify Chromatin IP system (#492024) were purchased from Thermo scientific. iScriptTM cDNA synthesis kit (#1708891), Clarity Max TM Western ECL substrate (#1705062), Clarity Western ECL substrate (#1705061), Immunoblot PVDF membrane (#1620177), iTaq TM Universal SYBRGreen super mix (#1725122) were procured from Bio-Rad. Glycine (#MB0131), TRIS (#MB-029), Sodium Dodecyl Sulphate (SDS) (#MB010), SM Powder (#GRM-1254), DMEM High Glucose (#AL151A), McCoy’s 5A (#AL057A), MEM (#AL047S), Glycerol (#GRM-081), Dimethyl sulphoxide (DMSO) (#TC185), BSA (#MB083) were procured from HiMedia. NFκB Activation Inhibitor II, JSH-23 (#CAS 749886-87-1) was purchased from Santacruz.

### 2.2 Cell culture

Human CRC cell lines-HCT116, HT29; human embryonic kidney cells HEK293 were procured from NCCS Pune. CT-26 mouse colon cancer cell line was a kind gift from Professor Sumit Kumar Hira, Cellular Immunology Laboratory, Department of Zoology, The University of Burdwan, Purba Bardhaman, India. The HCT116 cells were maintained in McCoy’s 5A, HT29 and CT-26 cells in DMEM, and HEK293 cells in MEM medium, added with 10% FBS and 1% antibiotic mixture. The cells were cultured in a CO_2_ incubator at 37[. The cells were seeded for experiments by detaching them using 0.05% Trypsin-EDTA. For *in vitro* studies cells were exposed to CDDP (30 µM), GSKJ4 (20 µM), or JSH23 (10 µM) at a confluency of 70%. CDDP was dissolved in 0.9% NaCl solution. GSKJ4 or JSH23 was added 1hr before CDDP exposure. Whenever required, NAC (20 mM) was administered 2 hrs prior to CDDP exposure. In the case of combination treatment with NAC, 1 hour after NAC administration, the inhibitors GSKJ4 or JSH23 were added. One hour after that, CDDP was added. All the treatment time points were 24 hrs except for the drug tolerant model.

### 2.3 DCFDA assay

For DCFDA microscopy analysis, 20 × 10^3^ cells were seeded on cover slides, while 2 × 10^5^ cells were seeded in 6 cm dishes for flow cytometry analysis. The cells were then kept in a CO_2_ incubator at 37[until it reached the desired confluency. Post-treatment, media was discarded followed by a PBS wash. After that, the cells were kept for 1 hr with 20 µM of DCFDA. The cells were then washed with 1X PBS (3 times with 1X PBS) solutions to remove excess DCFDA stain, then mounted with 70% glycerol and visualized under the microscope. For flow cytometric analysis, following DCFDA incubation, cells were washed with 1X PBS and treated with trypsin-EDTA to detach them. The cells were then collected and centrifuged at 2000 rpm for 3 minutes. The pellet was resuspended in 1 mL of 1X PBS, centrifuged again at 2000 rpm for 3 minutes, and finally resuspended in 500 µL of 1X PBS and DCFDA fluorescent intensity was analyzed through flow cytometry (Cytoflex, Beckmann Coulter).The DCFDA fluorimetry assay was performed following earlier reported protocol [47].

### 2.4 Cell viability assay

Cell viability assay was done using MTT. Approximately 6×10^3^ cells were cultured in a 96-well plate and allowed to grow overnight in the CO_2_ incubator. Subsequently, respective drug treatment was given for the stipulated period. After the incubation period was over, we added MTT (0.5mg/mL of media) to the desired wells and incubated for 3 hrs, followed by solubilizing the formazan crystal with DMSO. OD values were measured using a microplate spectrophotometer (Multiskan Sky, Thermo Scientific) at 570 nm and a differential filter of 630 nm.

### 2.5 Colony count assay using crystal violet

The crystal violet assay was performed by seeding 1×10^5^ cells in a 35 mm petri dish. After incubation, the media was aspirated, and the desired drug was added to the specific wells. After treatment, the drug containing media was discarded and a PBS wash was given. The cells were then fixed in chilled methanol for 20 mins and followed by a PBS wash. The cells were then subjected to 0.5% crystal violet solution and incubated for 10 mins, followed by rinsing with H_2_O. The cells were observed using an inverted microscope (10X).

### 2.6 Drug rescue experiment

2×10^5^ HCT116 cells were cultured in 6 cm plates. Post 24 hrs of CDDP exposure, the media was discarded from the petri plates and washed in PBS. Complete media was added, and the plates were kept for another 24 hrs in a CO_2_ incubator. For the drug tolerant model, after 3 days of CDDP exposure, the treatment media was aspirated, and the plates washed in 1X PBS. Then, fresh complete media was added, and incubation was performed for another 3 days.

### 2.7 Gene expression analysis

We isolated total RNA using Trizol reagent. cDNA synthesis was done using cDNA synthesis kit. We performed qRT-PCR using the SYBR Green PCR master mix in the QuantStudio 3 qRT-PCR system. The primers used are mentioned in Table 1. The mRNA expression of the genes was normalized to the housekeeping gene GAPDH or β-Actin.

**Table 1.**
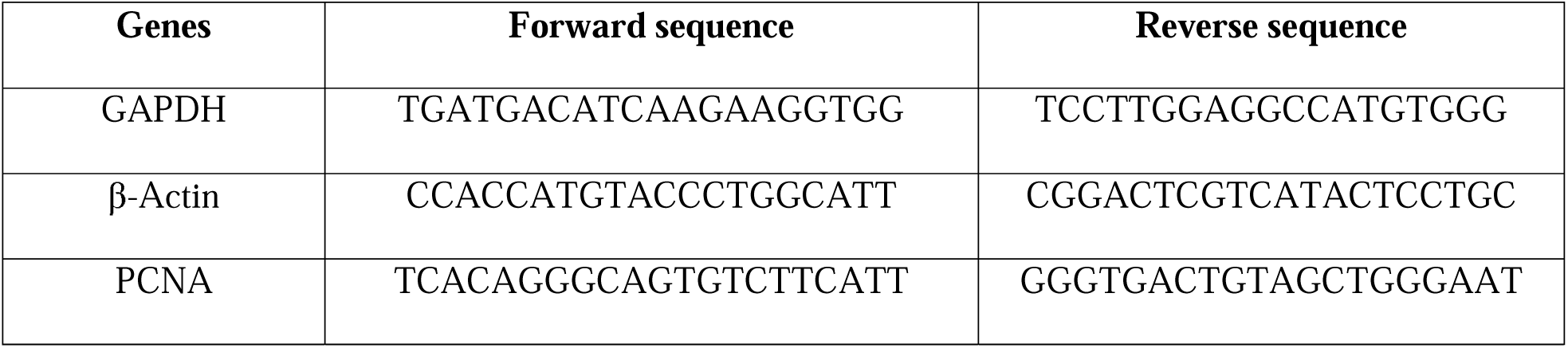
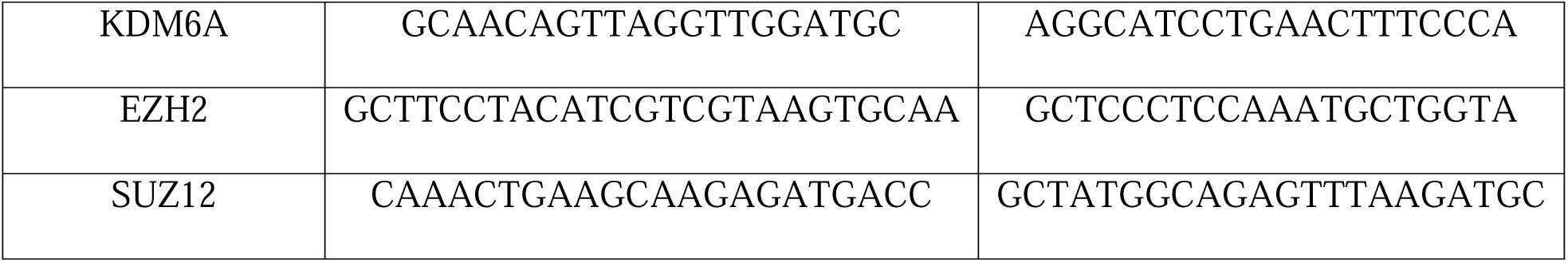
Primer sequences:

### 2.8 Immunoblotting

The total cellular protein lysate was extracted using RIPA buffer with added protease inhibitor, and quantified using Bradford reagent. An equal volume of protein samples was loaded into the well and resolved by sodium dodecyl sulfate polyacrylamide gel electrophoresis (SDS PAGE), followed by transfer to the PVDF membrane. The latter was then blocked by 5% skimmed milk made in 1X Tris buffered saline (TBS) and kept for 2 hrs at room temperature (RT). The membrane was then exposed to the primary antibody of interest at 4[overnight, followed by 2 hrs incubation with a secondary antibody (HRP-conjugated anti-rabbit or anti-mouse IgG). Finally, the bands were observed with ECL on ChemiDoc (Bio-Rad). β-Actin or GAPDH or Total ERK served as loading control for the immunoblot analysis. Protein levels were measured using ImageJ, normalized to loading control, and thereafter bar graphs were generated to represent comparative expression. A list of antibodies is mentioned in Table 2.

**Table 2:** List of antibodies:

| Antibody | Product details | Dilutions |
| --- | --- | --- |
| HP1 Alpha | Cell Signalling Technology # 2616 | 1:2000 |
| Tri-methyl-Histone H3 (Lys27) | Cell Signalling Technology (C36B11) # 9733 | 1:2000 |
| Histone H3 | Cell Signalling Technology (1B1B2) # 4269 | 1:3000 |
| EZH2 | Cell Signalling Technology (D2C9) # 5246 | 1:1000 |
| JARID2 | Cell Signalling Technology (D6M9X) # 13594 | 1:1000 |
| SUZ12 | Cell Signalling Technology (D39F6) # 3737 | 1:1000 |
| NF-κB P65 | Cell Signalling Technology (D14E12) # 8242 | 1:1000 |
| Phospho-NF-κB P65 (ser536) | Cell Signalling Technology (93H1) # 3033 | 1:1000 |
| PCNA | Cell Signalling Technology (D3H8P) # 13110 | 1:2000 |
| P21 | Cell Signalling Technology (12D1) # 2947 | 1:1000 |
| KDM6A/UTX | Cell Signalling Technology (D3Q1I) # 33510 | 1:1000 |
| CD44 | Cell Signalling Technology(156-3C11) # 3570 | 1:1000 |
| CD133 | Cell Signalling Technology (D2V8Q) # 64326 | 1:1000 |
| β-Actin | Cell Signalling Technology(8H10D10) # 3700 | 1:2000 |
| Anti-Rabbit IgG HRP-linked antibody | Anti-Rabbit IgG HRP linked antibody # 7074 | 1:3000 |
| Anti-Mouse IgG HRP-linked antibody | Anti-Rabbit IgG HRP linked antibody # 7076 | 1:5000 |

**Table 3.**
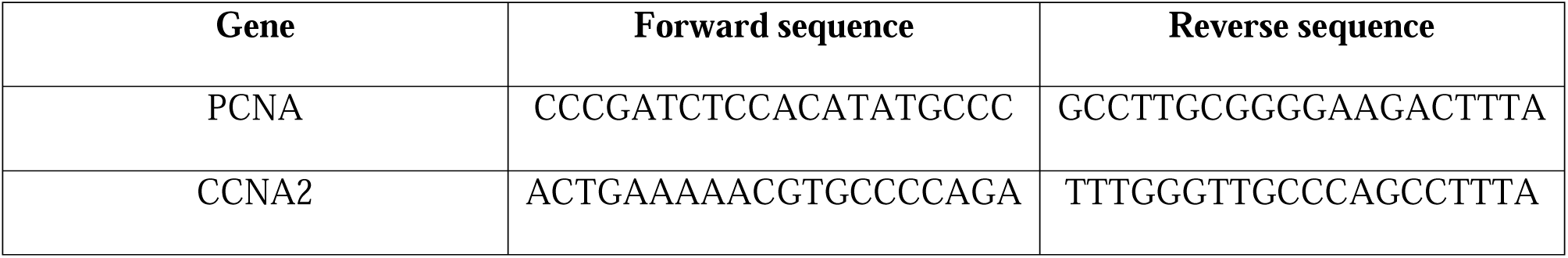
Promoter primer sequences:

### 2.9 Annexin V/Propidium Iodide (PI) apoptosis assay

2×10^5^ cells were seeded in a 6 cm dish and allowed to grow overnight. Post drug exposure, the treatment media was collected in a 15 ml centrifuge tube, the attached cells were trypsinized and collected in that tube as well, followed by centrifugation at 2000 rpm. Cell pellet was washed with 1X PBS at 2000 rpm, and the pellet was resuspended in 500 µl of 1X Annexin binding buffer. Annexin and propidium iodide (PI) were then added and incubated for 10 mins in the dark, except for the autofluorescence control. Cell death percentage was then measured in the flow cytometer (Cytoflex, Beckmann Coulter) using green filter FITC-A and red filter PE-A. 10000 events were captured, and the data analyzed using CytExpert software.

### 2.10 Cell cycle analysis

Post-treatment cells were trypsinized and pelleted by centrifugation at 2000 rpm. The pellet was thereafter resuspended in 100 µl of ice-cold PBS. To that 900 µl of chilled 70% ethanol (cell culture grade) was added dropwise. The cells were then kept at -20[overnight. The next day, cells were centrifuged at 3000 rpm, 4[, and then resuspended in PBS. Propidium iodide was then added to the cell suspension and incubated for 10 mins. Cell cycle phase distribution was analyzed through a flow cytometer. 10000 events were captured and analyzed through CytExpert.

### 2.11 Transient transfection

1×10^5^ cells per well were seeded in a 6 well plate and incubated overnight at 37[in a CO_2_ incubator. After that, to downregulate P65 or KDM6A, the cells were transfected using lipofectamine 3000 with siP65 (100 nM) (SignalSilence® NF-κB P65 siRNA II, Cell Signaling technology) or siKDM6A (50 nM) (IDT#hs.Ri.KDM6A.13.1), respectively. Scrambled RNA (40 nM) was used as a negative control.

### 2.12 Immunofluorescence microscopy

2×10^4^ cells were seeded on coverslips and allowed to grow till they reached 70% confluency. Post-treatment, coverslips were washed with PBS two times and fixed with 4% paraformaldehyde for 10 mins at RT. Cells were then permeabilized with 0.1% Triton X for 2 mins. The coverslips were then blocked with 2.5% BSA made in PBS for 1 hr. Further, cells were incubated overnight with the primary antibody of interest at 4[and the next day with corresponding secondary antibody. The nuclei were stained with DAPI for 10 mins. Finally, the coverslips were mounted in a clean glass slide using 70% glycerol. The cells were observed under a confocal microscope (Zeiss).

### 2.13 KDM6A/6B enzymatic activity assay

Nuclear proteins were harvested post-drug exposure using the standard nuclear protein extraction protocol [48]. To assess the KDM6A/6B enzyme activity, we used the KDM6A/6B Activity Quantification Assay Kit (# ab156910). We quantified the KDM6A/6B enzyme activity for its substrate in the CDDP-treated as well as untreated control groups, following the manufacturers protocol.

### 2.14 Drug-tolerant model

The drug-tolerant model was developed using HCT116 cells. To develop the model, we seeded 4×10^5^ cells in 10 cm dishes. We divided them into four distinct groups: a) untreated control, b) cells that received only CDDP (30 µM) for 3 days, c) cells that received 3 days of CDDP (30 µM) followed by 3 days of drug rescue, and d) cells that were subjected to CDDP (30 µM) for 3 days, followed by exposure to GSKJ4 (10 µM) for the next 3 days.

### 2.15 Chromatin immunoprecipitation assay (ChIP)

MAGnifyTM Chromatin Immunoprecipitation System was used to precipitate the soluble chromatin from the drug-treated samples with anti-EZH2, anti-KDM6A and a negative control anti-rabbit IgG. The total DNA considered as Input controls. Input controls and the ChIP samples were then amplified by qRT-PCR using the SYBR Green PCR master mix in the QuantStudio3 qRT-PCR system. The relative fold enrichment of our protein of interest has been quantified by normalizing with the input controls. The promoter primer sequences are mentioned in Table. 3.

### 2.16 Co-immunoprecipitation (Co-IP) assay

Total cell lysate was collected as described above. Protein G Magnetic Beads (HY-KO204-Med Chem Express) and IP-compatible anti-P65 and negative control anti-rabbit IgG have been incubated for 4 hrs in 4[using a rota spin. 500 µg of whole cell lysate was added to the respective tubes and incubated overnight at 4[using a rota spin. The next day, the antibody-bound protein complex was pulled and run through SDS PAGE along with its input controls. The interaction of P65 with our protein of interest was confirmed by probing it with specific primary and secondary antibodies.

### 2.17 *In vivo* syngeneic colon carcinoma model

Male BALB/c mice (6–8 weeks old, 25 ± 2 g body weight) were obtained from the central animal facility at BITS Pilani and studies were conducted with the approval of the Institutional Animal Ethics Committee (IAEC) at the Department of Pharmacy, BITS Pilani, Pilani campus (Protocol Number: IAEC/RES/33/06). The mice were subcutaneously injected in the hindquarter region with 5 × 10[CT-26 cells suspended in 100 µL of a 1:1 mixture of Matrigel and serum-free DMEM [49]. Ten days after tumor cell implantation upon the appearance of visible subcutaneous tumor mass, mice were randomized into groups. Tumor volume monitoring began on the day of randomization (considered as Day 0).

### 2.18 Tumor regression study

For the tumor regression study, mice were divided into three groups (n = 4 per group): (i) tumor-bearing vehicle control, (ii) CDDP-treated, and (iii) combination-treated with CDDP and GSKJ4. Treatment started on day 10 following CT-26 cell transplantation. Mice received a single dose of intraperitoneal (i.p.) injections of CDDP (20 mg/kg [50] and GSKJ4 (2 mg/kg) [51].

The CDDP solution was prepared by dissolving 0.5 mg of CDDP in a 1:1 mixture of 50 μL ethanol and 50 μL μL Tween 80, followed by the addition of 100 μL normal saline. For the combination treatment, 0.5 mg CDDP and 0.05 mg GSKJ4 were dissolved in the same solvent mixture (50 μL ethanol and 50 μL Tween 80), followed by 100 μL of normal saline. A total of 200 μL of this solution was administered per mouse [38]. Vehicle control mice received 200 μL of the solvent mixture (50 μL ethanol + 50 μL Tween 80 + 100 μL saline) *via* the i.p. route. Throughout the treatment period, animals were carefully monitored for tumor regression, locomotor activity, abnormal behaviour, and mortality. At the end of the treatment course on day 4, mice were euthanized, and the subcutaneous tumors were excised, sectioned, and subjected to further analysis.

### 2.19 Molecular docking studies

Docking of human Polycomb Repressive Complex 2 (PRC2) to the N-terminus region of human P65 protein was performed. The PRC2 complex, comprising Enhancer of Zeste Homolog 2 (EZH2), EED, SUZ12, and the cofactor JARID2 (PDB ID: 5HYN), was docked with the N-terminal region (residues 20– 314) of the human p65 protein (PDB ID: 1NFI, Chain A) using the ClusPro web server [52]. ClusPro employs a rigid-body docking algorithm and selects the optimal complex model based on a Fast Fourier transform (FFT) correlation approach.

### 2.20 Statistical analysis

Each experiment was repeated for a minimum of three times, and experimental values are expressed as mean ± SD. A comparison between the two groups was performed using the Student’s two-tailed t-test, and one-way ANOVA (Tukey) was used to compare more than two groups. All statistical analyses were done using GraphPad Prism 8.0.1. The data was considered significant if the p-value was ≤0.05. The symbols, *, **, ***, **** indicate p ≤0.05, p ≤0.01, p ≤0.001, p ≤0.0001 respectively.

## 3. Results

### 3.1 CDDP exposure results in an increase in histone repressive marks, EZH2 activity and induction of a non-proliferative state

Tumor cells have an astounding adaptive potential to diverse stress stimuli. To understand the adaptive chromatin modifications in CRC cells post-drug treatment, we exposed the cells (HCT116 and HT-29) to their respective IC50 doses of the FDA approved drug-cisplatin (CDDP) and analyzed the expression and localization of the different chromatin repressive tags. Interestingly, CDDP exposure resulted in a drastic increase in protein expression of the transcriptional repressive tag-Heterochromatin protein1-α (HP1-α) **(Fig. 1A)**. An evenly distributed localization of HP1-α was also observed in immunofluorescence staining **(Fig. 1B).** We then assessed the expression level of specific repressive marks H3K27me3 **(Fig. 1C & Supplementary Fig. 1A)**, and H3K9me3 in HCT116 and HT-29 cells as well **(Supplementary Fig. 1B)**. Interestingly, a drastic increase in protein expression of both H3K27me3 and H3K9me3 was evident. In corroboration to above, there was also a simultaneous increase in immunofluorescence for the above repressive tags. Notably, the localization of H3K27me3 was primarily restricted to the outer nuclear periphery compared to the evenly distributed localization of H3K9me3 post drug treatment **(Fig. 1D & Supplementary Fig. 1C-D).** Given that we observed a robust alteration in expression and unique localization of H3K27me3 after drug treatment, we were further interested to explore the effect of drug pressure on the family of proteins that are involved in catalysing the above repressive histone methylation- the PRC2 (Polycomb Repressive Complex 2). In spite of an overall increase in the histone repressive marks, indicating an attenuated transcriptional activity, the mRNA expression pattern of EZH2 followed a trend of increase **(Fig. 1E)**. Importantly, we also observed an elevated protein level of the key catalytic subunit-EZH2 **(Fig. 1F & Supplementary Fig. 1E)**, SUZ12 **(Supplementary Fig. 1F-G)**, and also JARID2 **(Supplementary Fig. 1H)** in the CDDP-treated cells compared to the control group in HCT116 and HT-29 cells. Furthermore, immunofluorescence staining of EZH2 also showed an increased nuclear localization of the protein in the CDDP-treated cells **(Fig. 1G).** These findings clearly suggest that CDDP exposure induces repressive epigenetic alterations in the CRC cells, which might be associated with their survival under drug stress. Interestingly, we observed that genes associated with cellular arrest or senescence like, P53 and P21 showed a parallel increase in their expression post drug stress indicating the acquisition of a non-proliferative state coupled to inculcation of repressive tags after drug exposure **(Fig. 1H-I)**. In accordance to above, flow cytometry analysis revealed a substantial increase in cells at the G2-M phase of cell cycle post CDDP treatment indicating an acquisition of a non-proliferative state **(Fig. 1J)**. Collectively, the above results indicate that tumor cells alter their epigenomic signature, especially induce repressive chromatin modifications coupled to growth arrest upon exposure to chemotherapeutic stress.

**Fig. 1.**
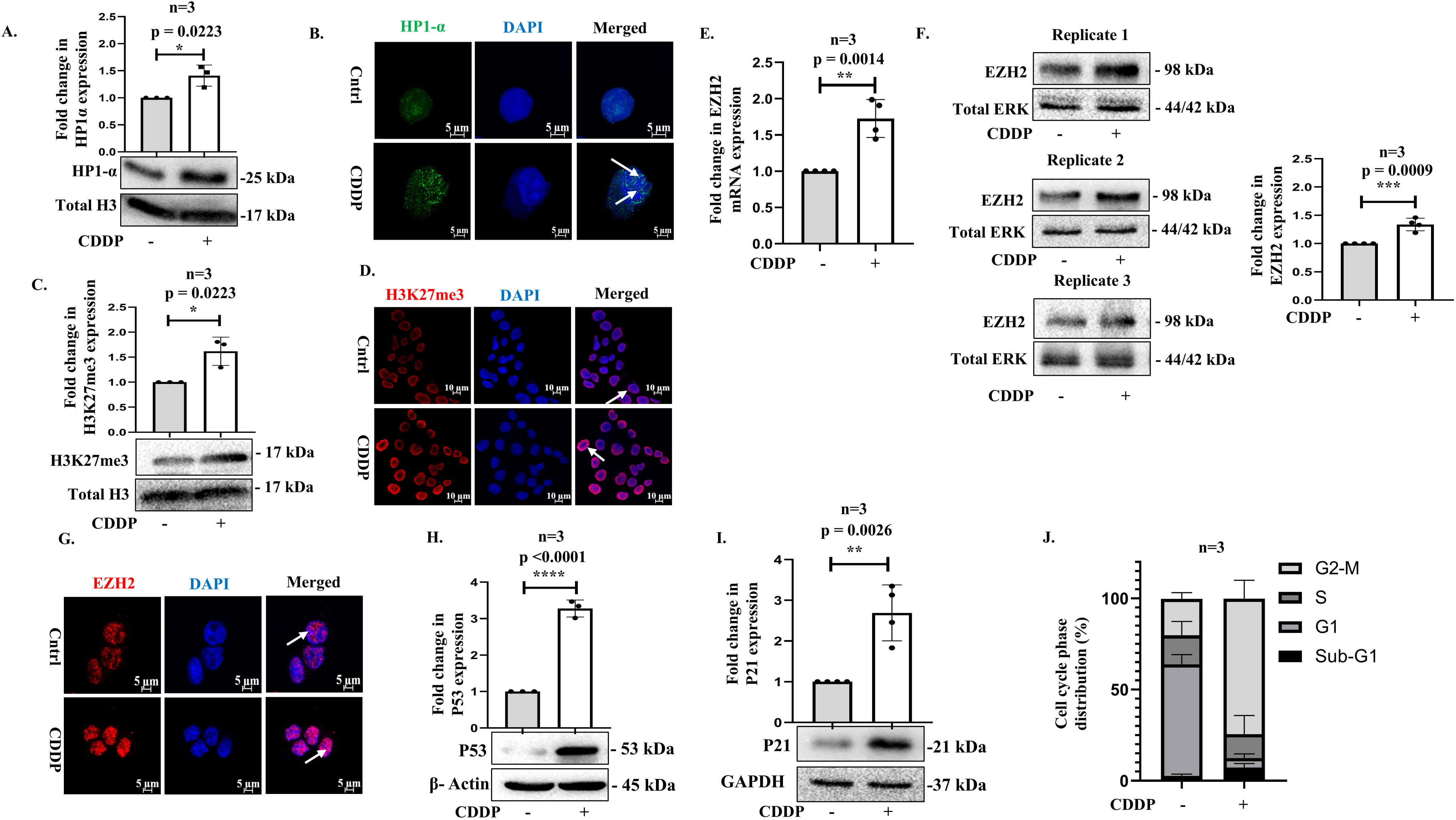
CDDP induces chromatin repressive marks and proliferative arrest: (**A**) Immunoblot and (**B**) immunofluorescence images showing expression and localization of HP1α (green) after CDDP (30 µM; 24 hrs). The cells were counterstained with DAPI (blue). Scale bar 5 µm. White arrows denote specific localization of the protein. (**C**) Expression of H3K27me3 as depicted through immunoblot; **(D)** immunofluorescence image showing localization of H3K27me3 (red). The cells were counterstained with DAPI. Scale bar 10 µm. **(E)** The mRNA expression of EZH2 as assessed by qRT-PCR analysis, followed by **(F)** immunoblots and **(G)** immunofluorescence staining (EZH2; red). Scale bar 5 µm. The cells were counterstained with DAPI. **(H-I)** The cellular expression levels of P53, P21 as assessed by immunoblots. **(J)** Cell cycle phase distribution analysis as performed by propidium iodide staining using a flow cytometer. Wherever applicable, CDDP treatment was at a dose of 30 µM for 24 hrs. Bar graphs show quantitative representation of the expression levels normalized to housekeeping gene or loading control. The data was considered significant (*) if the p-value was ≤0.05.

### 3.2 ROS regulates histone methyl transferase-EZH2-mediated induction of histone repressive marks at selected loci

Cancer cells usually generate abundant amount of ROS to aid their need for migration, metastasis, and enhanced proliferation. Conversely, increasing ROS level has been utilized as an approach to tip the balance toward cell death [53]. However, the role of ROS as a trigger controlling adaptive epigenetic modifications is poorly explored. Interestingly, fluorimetry, flow cytometric, and microscopic image analysis convincingly demonstrated that there was a significant induction of intracellular ROS after exposure of CDDP to HCT116 **(Fig. 2A & Supplementary Fig. 2A-B)**, while a marginal increase was evident in HT-29 cells **(Supplementary Fig. 2C)**. Importantly, quenching of intracellular ROS using N-acetyl-cysteine (NAC) resulted in a partial rescue of CDDP cytotoxicity suggesting that oxidative stress is responsible for the cytotoxic effect and therefore can be a key player modulating the epigenetic adaptive mechanisms acquired under drug stress **(Supplementary Fig. 2D-E).** In accordance to above, quenching of ROS resulted in a reversible alteration of the chromatin repressive state, as evident from the substantial reduction in expression of H3K27me3 marks **(Fig. 2B-C)** and PRC2 proteins such as EZH2 **(Fig. 2D-E)**, SUZ12 (**Fig. 2F)** and JARID2 **(Supplementary Fig. 2F).** A reversal of chromatin repression in CRC cells was also associated with a concomitant downregulation of P21 **(Fig. 2G)** and an increased expression of the proliferating cell nuclear antigen (PCNA) levels (both mRNA and protein level), which is known to be involved in DNA replication, indicating a reacquisition of the proliferative potential by the CRC cells **(Fig. 2H-J)**. Herein, we hypothesized that drug stress drives the cells towards a transcriptional shutdown and a coupled non-proliferative state mediated by preferential binding of EZH2 to the upstream promoter elements of proliferation associated genes. Importantly, chromatin immunoprecipitation (ChIP) assay showed a significant enrichment of EZH2 at the upstream promoter region of both PCNA and CCNA2 under drug stress. In contrast, NAC-treated cells showed a reduced promoter occupancy of EZH2 on the above genetic elements **(Fig. 2K-L)**. These data clearly indicate that EZH2 plays a critical role in regulating chromatin status through selective binding at genetic loci which might be putatively involved with a transitory non-proliferative and transcriptionally repressive state of the CRC cells under cisplatin stress.

**Fig. 2.**
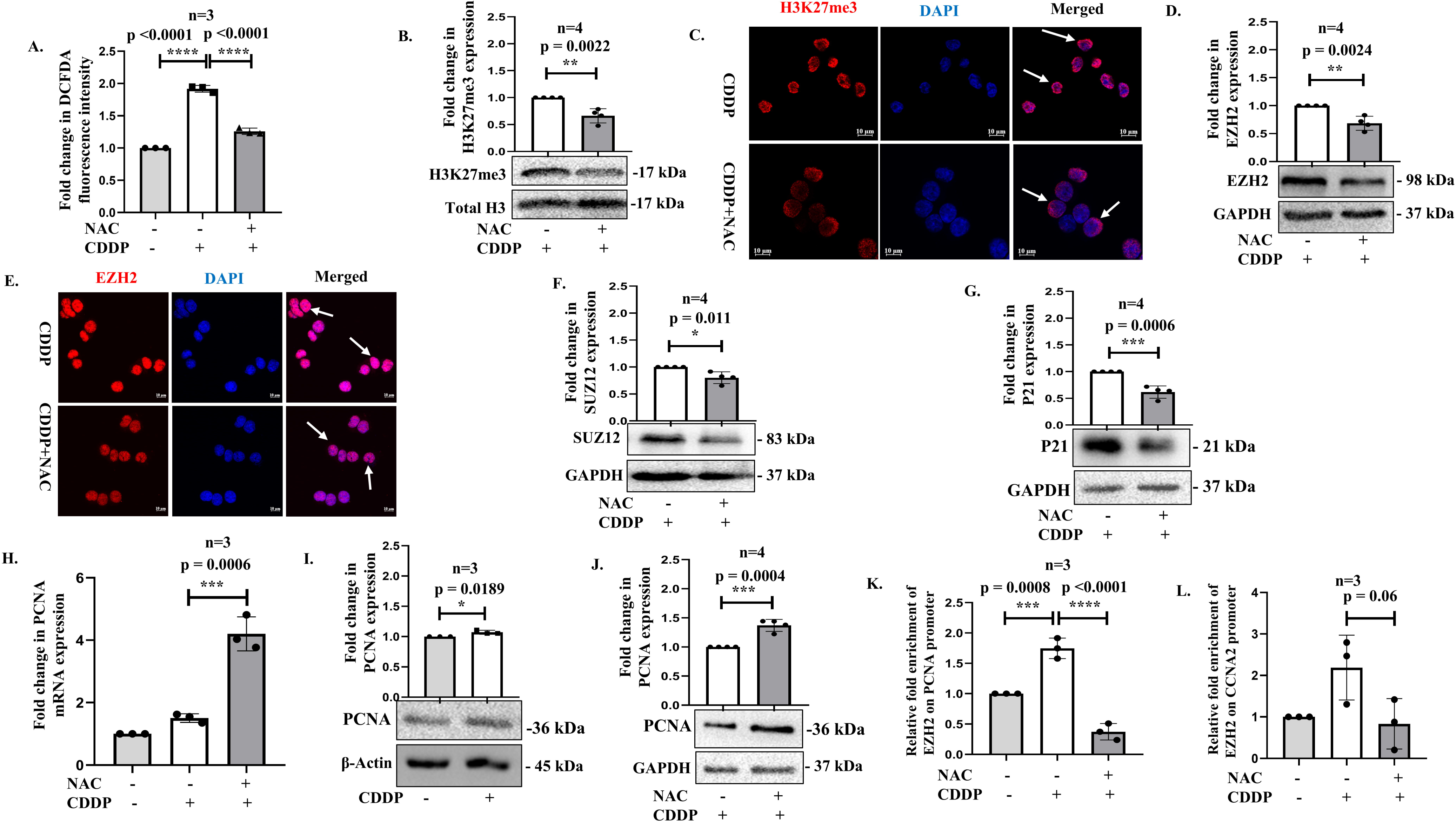
Quenching of ROS results in reversal of chromatin repressive marks and promoter occupancy of EZH2: **(A)** Bar graph showing relative DCFDA fluorescence intensity in cisplatin-treated samples with or without NAC (20 mM). **(B)** Immunoblot and **(C)** immunofluorescence images of H3K27me3 (red) after cisplatin treatment with or without NAC. The cells were counterstained with DAPI. Scale bar 10 μm. White arrows denote specific localization of the protein. **(D)** Immunoblot and **(E)** immunofluorescence analysis showing EZH2 (red) expression with or without NAC. The cells were counterstained with DAPI. Scale bar 10 μm. **(F)** SUZ12 expression as analyzed through immunoblot**. (G)** Effect of NAC on P21 protein expression as measured through immunoblot. Bar graph and representative immunoblots respectively showing the mRNA (**H**) and protein expression (**I-J**) of PCNA before and after CDDP treatment, as well as CDDP in combination with NAC. (**K-L**) ChIP assay demonstrating relative fold enrichment of EZH2 on PCNA and CCNA2 upstream promoter element. Wherever applicable, CDDP treatment was at a dose of 30 µM for 24 hrs. Bar graphs show quantitative representation of the expression levels normalized to housekeeping gene or loading control. The data was considered significant (*) if the p-value was ≤0.05.

### 3.3 NF-κB-P65 regulates and interacts with EZH2 governing its function

ROS involves generation of peroxides and superoxide radicals in the tumor cells which in turn has the potential to trigger specific cellular signaling pathways that can transduce diverse effects at the epigenetic level as well [54–56]. Thus, having established the involvement of ROS in epigenetic dysregulation and associated attenuation of proliferative gene transcription, we were interested to analyze the molecular signaling cascades that have a key role downstream of ROS- in this regard existing reports emphasize the role of the P65 component of the NF-KB signaling pathway. Importantly, we observed a dramatically increased expression of phospho-NF-KB P65 (ser536) after drug exposure and quenching of ROS resulted in a reduction in its expression level **(Fig. 3A-B)**. Moreover, nuclear localization of the phospho-NF-KB P65 subunit significantly increased after CDDP exposure **(Fig. 3C)** indicating a putative nuclear function; however, there was a reversal of expression and localization when ROS levels were ablated using NAC **(Supplementary Fig. 3A)**. The above experiments imply the role of P65 in regulating downstream function of ROS post CDDP treatment. We hypothesized that P65 can be possibly involved in modulation of the epigenome, as a significant alteration of epigenetic signature was simultaneously observed. Interestingly, pharmacological inhibition (using JSH23) or genetic ablation of P65 (siRNA) caused a reduction in p-P65 and EZH2 protein levels and its nuclear localization **(Fig. 3D-G & Supplementary Fig. 3B)**, alongside a decrease in the senescence marker P21 **(Fig. 3H-I)**. The above findings imply that P65 is upstream to EZH2 and its activity after drug stress, however, we were interested to explore the role of P65 inside the nucleus as well. Interestingly, co-immunoprecipitation study showed that P65 physically interacts with EZH2 and thus can be critical to EZH2-mediated inculcation of repressive tags post drug stress **(Fig. 3J)**. Earlier, reports have shown that P65 can interact with EZH2 but in a completely different context [57,58]; ours is the first study to show their interaction under drug stress. Taken together, the above results suggest that NF-KB-P65 regulates the activity and expression of EZH2 which in turn facilitates chromatin repression through H3K27me3 accumulation. Interestingly, we observed that not only EZH2, P65 also regulates the expression of another PRC2 family protein SUZ12 both at gene and protein level **(Supplementary Fig. 3C-D)**. In accordance to the previous findings, both siRNA-mediated knockdown of P65 and its pharmacological inhibition led to a reduction in SUZ12 mRNA and protein expression. In contrast, JARID2 expression remained comparable to that observed in the CDDP-treated group **(Supplementary Fig. 3E)**. Furthermore, co-immunoprecipitation assay revealed that P65 interacts with SUZ12 and interestingly, pharmacological inhibition of P65 led to a marked reduction in the interaction between these two proteins **(Supplementary Fig. 3F)**. Moreover, to understand the plausible interactions between P65 protein with the members of the PRC2 complex, we performed molecular docking studies between P65 and PRC2 complex containing EZH2, SUZ12, EED and JARID2 proteins. Docking analysis proposes a possible interaction between P65 and core members of the PRC2 complex including EZH2, SUZ12 and EED. Three distinct poses from the top five docking results **(Fig. 3K)** were superimposed onto the X-ray structure of the PRC2 complex (PDB ID: 5HYN). The docking analysis suggests that the N-terminal domain of P65 can interact with the PRC2 complex *via* multiple regions. In the first pose (green ribbon), P65 approaches the interface where the EED protein binds to EZH2, making contacts with both EED and EZH2. In the second pose (yellow), P65 interacts near the region where SUZ12 is bound to EZH2. In the third pose (purple), P65 binds exclusively to EZH2, without contacting EED or SUZ12.

**Fig. 3.**
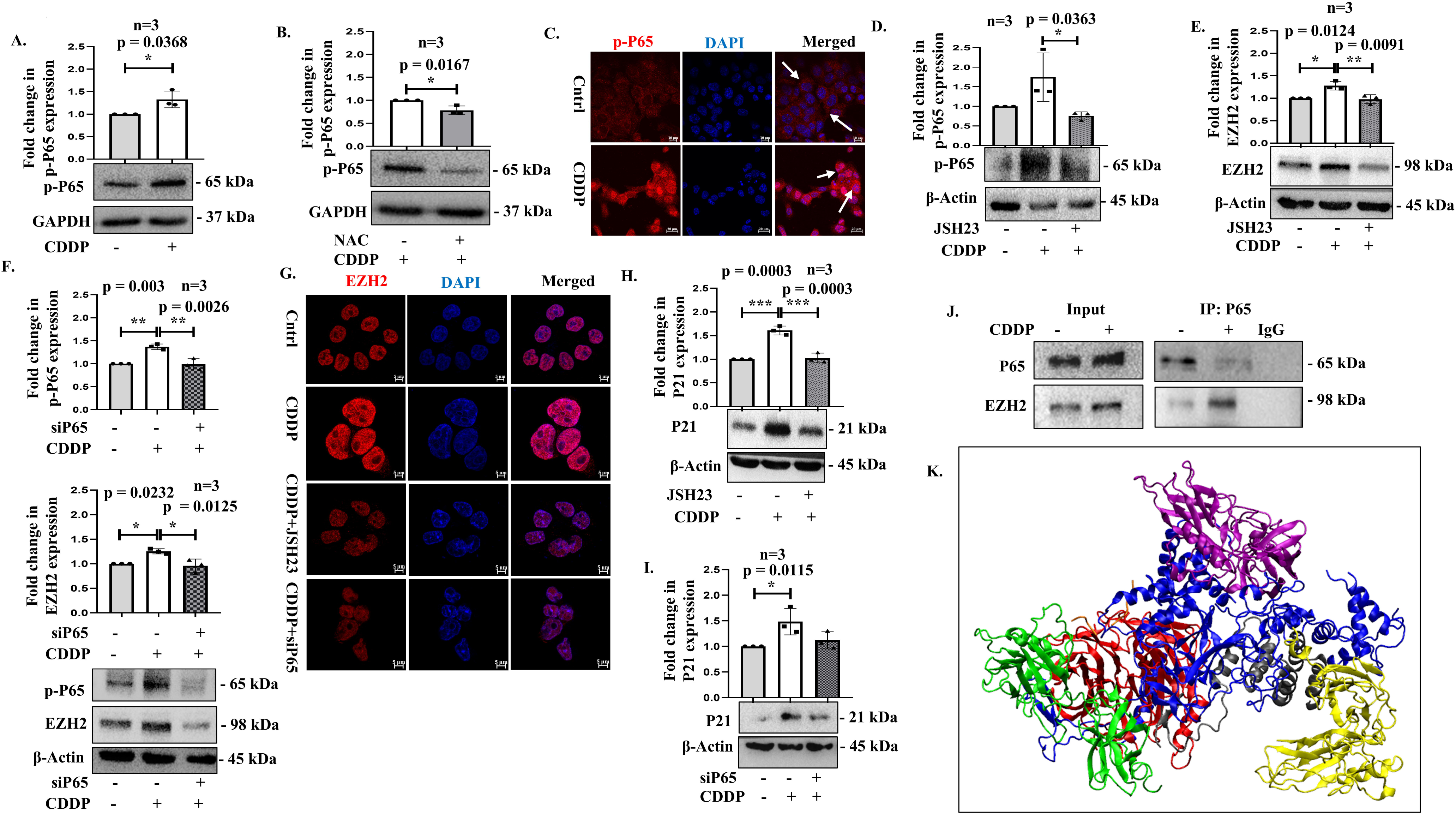
ROS-induced P65 regulates EZH2 expression: **(A)** Bar graph showing the protein expression of p-P65 in the CDDP-treated cells compared to untreated control. **(B)** Immunoblot depicting change in expression of p-P65 in the NAC-treated cells compared to CDDP. **(C)** Localization of p-P65 (red) post CDDP exposure as depicted through immunofluorescence images. The cells were counterstained with DAPI (blue). Scale bar 10 μm. White arrows denote specific localization of the protein. **(D-F)** Immunoblots showing expression of p-P65 and EZH2 after treatment with CDDP with or without JSH23 or siP65. **(G)** Cellular localization of EZH2 (red) as assessed through immunofluorescence images. The cells were counterstained with DAPI (blue). Scale bar 5 μm. **(H-I)** Immunoblots showing expression of P21 post P65 inhibition (JSH23 or siP65). **(J)** Co-immunoprecipitation assay representing interaction between P65 and EZH2. Immunoprecipitation was performed with anti-P65 antibody and probed with desired antibody. Total cellular lysate served as input. (**K**) Molecular docking of the human P65 protein (PDB ID: 1NFI) with the PRC2 complex (PDB ID: 5HYN): A ribbon diagram depicting the top three distinct docking poses of the P65 protein (green, purple, and orange ribbon) is shown superimposed on the PRC2 complex. The components of the PRC2 complex-EZH2, EED, SUZ12, and JARID2 are represented by blue, red, black, and orange ribbons, respectively. Wherever applicable, CDDP treatment was at a dose of 30 µM for 24 hrs. Bar graphs show quantitative representation of the expression levels normalized to housekeeping gene or loading control. The data was considered significant (*) if the p-value was ≤0.05.

### 3.4 ROS inhibition or drug withdrawal induces histone demethylase-KDM6A activity which in turn re-initiates cellular proliferation

The histone demethylases are a family of proteins with Jumonji C (JmjC) domains that act antagonistic to histone methyltransferases with potential to remove methylation marks from lysine/arginine residues of histone proteins utilizing Fe(II) and alpha-ketoglutarate (αKG) as cofactors [59,60]. These JmjC-containing proteins are already implicated in several cancers. Herein, UTX/KDM6A is especially responsible for removal of H3K27me2/3 marks. This enzyme has been shown to be associated with loss-of-function mutations and is believed to be a tumor suppressor in multiple cancers like renal carcinoma, bladder cancer and leukemia [61–64]. In contrast, it is also reported to be frequently overexpressed in several other cancers types like lung and breast cancer thus playing a pro-tumorigenic function as well [65,66]; however its role in CRC is poorly explored. Given that we observed a spontaneous increase in H3K27me3 mark upon drug stress, and its dynamic reversal upon quenching of ROS motivated us to analyse the role of the H3K27me3 demethylase (KDM6A) under the above conditions. Importantly, an elevated mRNA and protein expression of KDM6A were observed in cells upon ROS quenching implicating its probable role **(Supplementary Fig. 4A & Fig. 4A)**. To negate a specific NAC-dependent effect, we used an alternate ROS scavenger as well-ascorbic acid and found a similar trend in cellular proliferation as well as KDM6A protein expression **(Supplementary Fig. 4B-C)**. An increase in KDM6A and a decrease in H3K27me3 do not always imply a probable direct role of KDM6A, as it can be resultant of reduced EZH2 function as well. Therefore, to prove the same we analyzed the enzymatic activity of KDM6A which showed an increase in the NAC-treated group compared to CDDP **(Fig. 4B)**. Importantly, similar results with increased KDM6A expression were also observed in CRC cells after a drug-rescue or drug vacation **(Fig. 4C),** which correlates with the reduction in H3K27me3 protein level **(Fig. 4D)**. Herein, we hypothesized that the reversal of tumor cells from the drug-induced growth-arrest state to a proliferative one upon ROS quenching or drug withdrawal is associated with a switch in activation of KDM6A resulting in chromatin modulation conducive to re-emergence of the proliferative ability or PCNA expression of the CRC cells **(Fig. 4E).** To delineate the role of KDM6A in this context, we performed a ChIP assay. Interestingly, KDM6A enrichment was now observed on upstream promoter elements of proliferation associated genes like PCNA and CCNA2 in the NAC-treated groups compared to the CDDP-treated cells and untreated control **(Fig. 4F-G)**. We, thereafter checked the protein-protein interaction of PCNA and KDM6A through immunoprecipitation assay and observed that they interact with each other as well **(Supplementary Fig. 4D)**. This suggests that KDM6A actively regulates cancer cell proliferation by dynamically eliminating the di or trimethylated H3K27 repressive marks from specific genes involved in cellular proliferation and also regulates gene expression through protein-protein interaction as well. Overall, herein, we established an epigenetic switch between EZH2 and KDM6A in CRC cells, which regulates the active or repressed state of specific genes in a context-dependent manner.

**Fig. 4.**
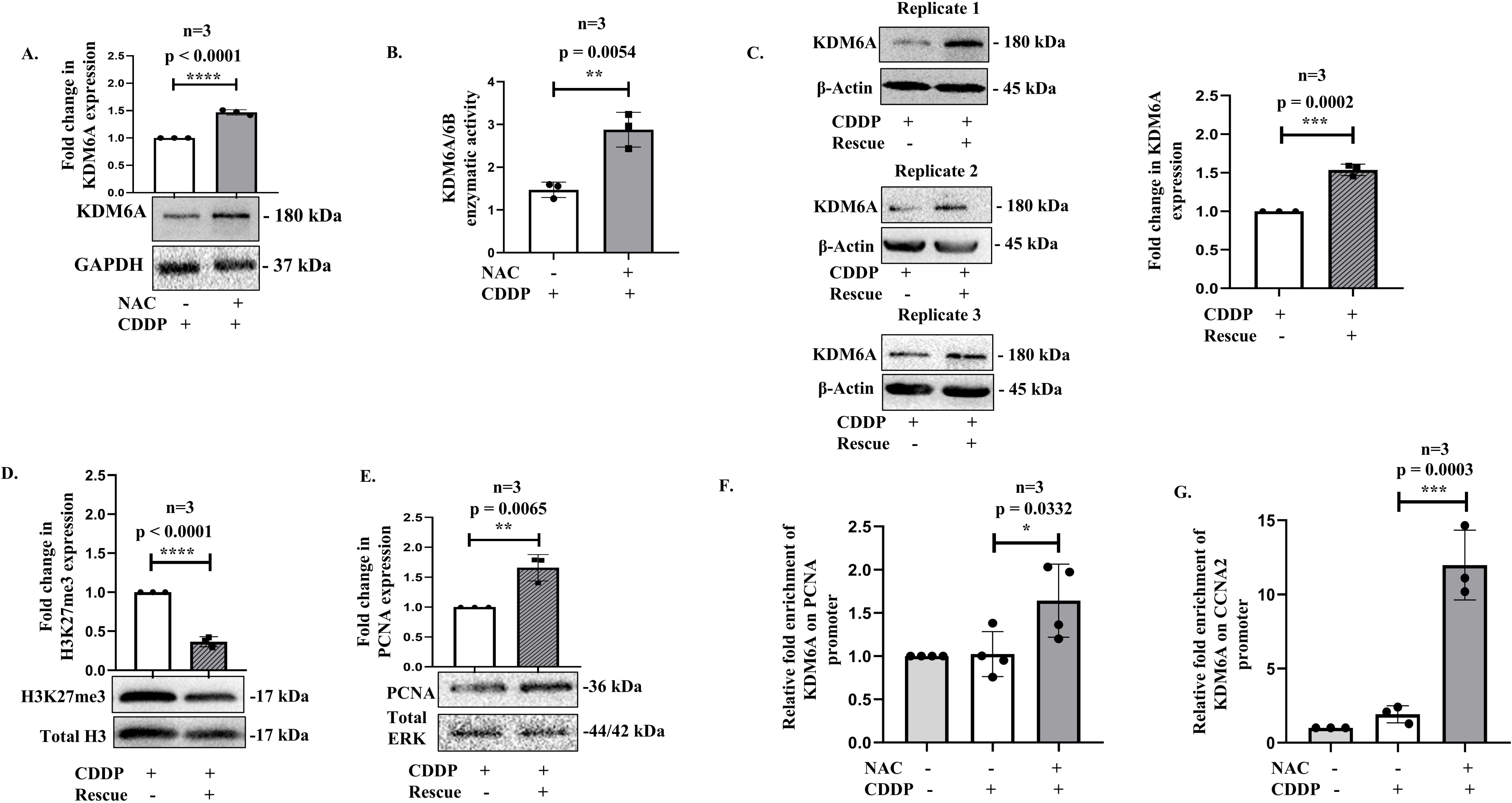
Quenching of ROS or drug withdrawal results in increased activity of KDM6A associated with cellular proliferation: **(A)** KDM6A protein level after NAC treatment compared to only CDDP as analysed through immunoblots. **(B)** The enzymatic activity of KDM6A/6B after CDDP exposure in presence or absence of NAC. **(C-E)** Protein expression of KDM6A, H3K27me3 and PCNA. Cells were treated with CDDP (30 μM; 24 hrs) followed by withdrawal of the drug from culture media and further incubation for another 24 hrs. **(F-G)** Bar graph showing relative fold enrichment of KDM6A on PCNA and CCNA2 upstream promoter region after CDDP exposure with or without NAC compared to untreated control. Wherever applicable, CDDP treatment was at a dose of 30 µM for 24 hrs. Bar graphs show quantitative representation of the expression levels normalized to housekeeping gene or loading control. The data was considered significant (*) if the p-value was ≤0.05.

### 3.5 Inhibition of KDM6A activity as a potent strategy to prevent tumor growth and relapse after chemotherapy

Based on the above results, it is evident that KDM6A can play a pivotal role in the proliferative ability of the CRC cells and the demethylase can hence be a suitable therapeutic target to prevent CRC relapse. Therefore, to validate the above, we inhibited KDM6A both pharmacologically-with a potent small molecule inhibitor (GSK-J4), and also *via* genetic manipulation (siKDM6A). As expected, inhibition of KDM6A **(Supplementary Fig. 5A-B)** resulted in a considerable increase in H3K27me3 **(Supplementary Fig. 5C-D)**. Importantly, KDM6A inhibition resulted in a reduced expression of PCNA **(Fig. 5A-C)** and this was also associated with a significant cell death as analyzed by Annexin V/PI based flow cytometry **(Supplementary Fig. 5E)**. Importantly, a higher percentage of cell death was observed in CDDP plus GSKJ4 samples compared to only CDDP, and in CDDP+NAC+GSKJ4-treated groups in comparison to CDDP plus NAC. This motivated us to explore the potential effects of GSKJ4 on drug-tolerant persister (DTP) cells that survive CDDP pressure and contribute to the re-initiation of proliferation overtime, if given a drug vacation. To simulate the above putative clinical condition, CRC cells were treated with GSKJ4 (3 days) post CDDP exposure (3 days) or given a 3-day drug holiday after CDDP exposure **(Supplementary Fig. 5F).** Interestingly, cells treated with GSKJ4 and CDDP showed a drastically reduced cellular revival when compared to only CDDP-treated cells **(Fig. 5D)**. Immunoblot analysis also showed an elevated level of KDM6A and PCNA in the CDDP-rescue group only, and not in GSKJ4, suggesting the role of KDM6A in revival post drug stress **(Fig. 5E-F)**. The above results highlight the importance of KDM6A in re-emergence of proliferative cells after drug treatment. Herein, several reports suggest that KDM6A can regulate the expression of cancer stemness which is often associated with enhanced cellular proliferation and aggressive cellular behaviour [67,68]. In corroboration to above, we observed an increased expression of CD44 and CD133 in the CDDP-rescue group but not in cells exposed to GSKJ4 alongside CDDP. This clearly indicates that KDM6A inhibition can restrict revival of CRC cells after cisplatin therapy **(Fig. 5G-H)**. Similar results were observed when KDM6A was inhibited in CDDP treated HCT116 cells, along with NAC for 24hrs, where both CD44 and CD133 expression levels were reduced **(Supplementary Fig. 5G-H)**. To investigate the potential correlations between KDM6A and tumor growth, we performed an *in vivo* tumor regression study using a CT-26 mouse cell line derived syngeneic colon carcinoma model in BALB/c mice. Interestingly, aligned to our *in vitro* findings, CDDP plus GSKJ4 treated mice showed a significantly increased tumor regression compared to only CDDP-treated mice **(Fig. 5I-J)**. This was associated with a reduction in expression of stem cell associated markers like CD44 in the KDM6A inhibited mice **(Supplementary Fig. 5I).** Importantly, to translate KDM6A inhibitors from bench to bedside they must not show cytotoxic effect on non-tumor cells; and importantly, KDM6A inhibition in human embryonic kidney (HEK293) cells had no effect on their proliferative ability providing a platform for futuristic studies with KDM6A inhibitors **(Supplementary Fig. 6A-D**).

**Fig. 5.**
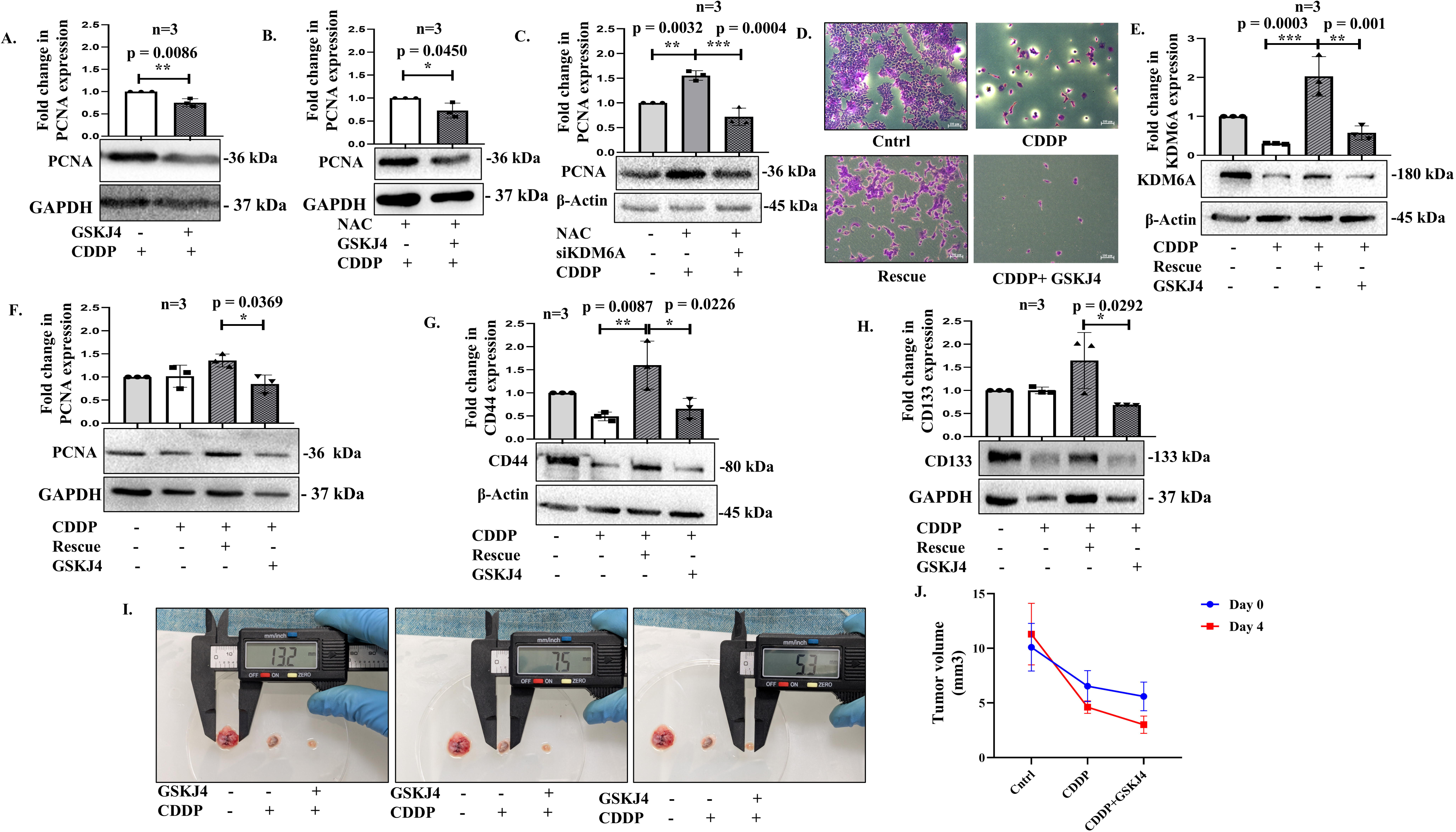
Inhibition of KDM6A prevents re-proliferation of tumor cells post CDDP treatment: (A-C) Expression level of PCNA post-KDM6A inhibition through GSKJ4 or siKDM6A knockdown. CDDP treatment was at a dose of 30 µM for 24 hrs. **(D)** Crystal violet images comparing the cell viability in all the above-mentioned experimental groups**. (E-F)** Immunoblots showing expression of KDM6A and PCNA in the drug-tolerant cells under drug rescue or KDM6A inhibition. **(G-H)** Expression of CD44 and CD133 as observed through immunoblots in the drug tolerant cells. **(I)** Representative images of tumors collected on days 0, 4, and 8 following the initiation of treatment highlighting differences in tumor regression among the treatment groups: vehicle control, CDDP, and the combination of CDDP with GSKJ4. **(J)** The graph illustrates the comparative reduction in tumor volume among the treatment groups between day 0 (treatment initiation) and day 4 (treatment termination). The data was considered significant (*) if the p-value was ≤0.05.

## 4. Discussion

Existing studies outline an observed phenomenon in cancer therapy called “retreatment response” or sensitivity after a “drug holiday.” For example, non-small cell lung cancer patients who earlier responded to EGFR inhibitors later on experienced therapy failure due to resistance, but the same patients demonstrated a subsequent response to the inhibitors post a “drug vacation”[4,69]. This suggested that resistance to chemotherapy might entail a reversible phase, which is probably unlikely due to stable genetic alterations. In this regard, several comprehensive molecular studies have revealed that different chromatin modifying molecules are frequently altered in cancers and facilitate drug tolerance [70,71]. In this regard, EZH2 has been found to be intricately associated with tumor stemness and shorter survival of cancer patients, including in colorectal/gastrointestinal cancers. These studies primarily describe the consequences of EZH2 mutations or over-expression positively contributing to tumor metastasis or aggressiveness. In our study, we observed an increase in EZH2 after drug stress, which we believe is kind of an adaptive stress-response strategy of the tumor cells to survive the drug pressure, thus playing a positive and crucial role in sustenance of tumor cells eventually contributing to relapse. Herein, a few existing reports also support this role of EZH2 in adaptive resistance, and regulation of tumor cell survival under drug stress [72–74]. This compensatory epigenetic survival response adds to the well-established role of EZH2 in enhancing tumor aggressiveness as discussed above [75]. Further, in our study, an inhibition of EZH2 with GSK126, which binds to the S-adenosylmethionine (SAM) binding pocket of EZH2 and shows competitive inhibition SAM, was found to reduce the histone repressive marks post cisplatin exposure suggesting that EZH2-mediated adaptive stress response is an outcome of its core catalytic activity. In corroboration to above, we observed epigenetic alterations facilitating the transient dormant state in CRC cells, marked by an up-regulation of EZH2, and a coupled chromatin repression imbibing survival adaptive response in CRC cells post-CDDP exposure. While a myriad of reports suggest the involvement of EZH2 in cancer proliferation; enable tolerance towards glucose deprivation [76]; or suppress ferroptosis reducing sorafenib sensitivity in HCC [77], its role in modulating cisplatin sensitivity in CRC cells is unique [78]. Importantly, our study also portrays the fundamental role of ROS as an upstream regulator of EZH2 and in dictating epigenetic status of CRC cells. NAC-mediated ROS quenching reversed EZH2-induced chromatin re-modelling. Interestingly, the P65 component of the NF-KB signaling was activated by ROS and its inhibition resulted in decreased EZH2 levels, revealing a key link between redox response and epigenome. An earlier report suggests that P65 can regulate transcription of EZH2 by binding to its intronic elements [79]; however, we found that the two proteins physically interact and can regulate drug response.

While EZH2 catalyses methylation of histone residues, its activity is countered by the histone demethylases (HDMs), consisting of a family of proteins namely KDMs. Similar to histone methyltransferases (HMTs) these proteins are also implicated in therapy resistance and treatment failure in multiple cancer types [80,81]. Furthermore, interestingly, KDM6A has been controversially discussed in literature with respect to cancer. While a substantial number of reports portray KDM6A as a potent tumor suppressor in various gastrointestinal cancers [82,83], including colorectal cancer (CRC); however, several studies have also reported the pro-tumorigenic role of KDM6A, particularly through its influence on chromatin remodelling and transcriptional regulation of oncogenic pathways. In favour of its pro-tumorigenic function, for example, Zhang *et al.* 2020 have shown that inhibition of KDM6A can repress the tumor-initiating cells and stemness related genes [84]. Furthermore, Wang *et al.* 2020 reported that KDM6A inhibitor GSKJ4 along with oxaliplatin, can dramatically reduce the load of colorectal cancer [85]. We therefore assume that the role of KDM6A is purely context-dependent and might depend on tumor type, stage, therapy procedure and on interacting proteins as well. We report a trigger in expression of KDM6A when given a drug interval, or when ROS is quenched leading to activation of genes associated with re-initiation of tumor growth. Further, in our study, KDM6A was found to occupy the upstream elements of growth promoting genes, under the above-mentioned conditions thus establishing its role in selective chromatin regulation. Interestingly, Sharma *et al* in 2010 showed that drug tolerant cells survive through IGF-1 receptor signaling and a modified chromatin state maintained by another type of histone demethylase-KDM5A [4]. Truly, the KDMs constitute a large family of proteins, with diverse substrates and context dependent functions, thus demanding further exploration of their role. Herein, in our study, we found that a dynamic balance between EZH2 and KDM6A modulates the chromatin and associated gene expression post drug stress; a high EZH2 level reinforced chromatin repression, while a switch to elevated KDM6A activity corresponded to a more open chromatin state and re-initiation of proliferation ability of the CRC cells. Importantly, KDM6A is already reported to be associated with CBP to mediate permissive chromatin state on estrogen receptor (ER) targets promoting breast cancer (BC) progression [86]. KDM6A and MLL were also found to cooperatively mediate BC cell proliferation through creation of open chromatin marks [87]. In accordance to above, targeting KDM6A using pharmacological inhibitor-GSKJ4 or siKDM6A not only resulted in a pronounced cytotoxicity but also curbed the potency of long-term drug-treated CRC cells to reinstate their proliferative ability suggesting that KDM6A is not merely a bystander but a crucial player in dictating tumor relapse.

In recent times, epigenetic reprogramming has gained limelight as a promising strategy to eradicate drug-tolerant cancer cells [88,89]. Despite the DNA methyl-transferase inhibitor’s FDA approval, their poor pharmacokinetics represent a major setback [90]; further, HDACi are also approved, but they only target ∼10% of all acetylation sites [91]. Hence, efforts are currently being undertaken to discover new molecules that can be more effective and selectively inhibit specific epigenetic adaptations. In this regard, a shift in focus to targeting specific KDMs might be promising. Although much remains to be explored, especially understanding the efficacy of KDM6A inhibition in patient derived culture samples, this study however opens up an epigenetic basis of relapse and therefore a prospective novel therapeutic avenue. Our findings may not fully capture the complexity of tumor heterogeneity, microenvironmental influences, and immune interactions that exist in actual patient tumors. Therefore, future studies involving clinical samples and xenograft models are essential to evaluate the translational relevance and therapeutic efficacy of targeting KDM6A or EZH2 in CRC relapse.

## 5. Conclusions

Cells that maintain viability after an acute drug-shock represent the pool of cells that persist to contribute to recurrence. Interestingly, we observed that the persister cells undergo an overhaul of epigenome. A distinct chromatin-state maintained by KDM6A and EZH2 is pivotal to establishment of the minimal-growth yet pro-survival chromatin or revival of proliferative state. Our study unravels the transcriptional-regulatory activity of KDM6A and EZH2 which provide a novel understanding of how multi-domain transcriptional regulators control gene expression under drug-shock.

## List of Abbreviations

CRC: colorectal cancer
CDDP: cis-diamminedichloroplatinum (II)
PBS: Phosphate Buffer Saline
PRC2: Polycomb Repressive Complex 2
EZH2: Enhancer of Zeste homolog 2
KDM6A: Lysine demethylase 6A
ROS: Reactive oxygen species
NAC: N-acetyl-L-cysteine
DCFDA: 2′,7′- dichlorofuorescin diacetate
PCNA: Proliferating cell nuclear antigen
ChIP: Chromatin Immunoprecipitation
Co-IP: Co-Immunoprecipitation
ECL: Enhanced Chemiluminescence
PVDF: Polyvinylidene fluoride
RIPA buffer: Radioimmunoprecipitation assay buffer
PI: Protease inhibitor
i.p.: Intraperitoneal

## Declarations

### Consent for publication

All authors read and approved the final manuscript.

### Availability of data and materials

The datasets used and/or analyzed during the current study are available as a part of results – discussion section as well as in the, supplementary information files. All original data sets and replicates are also uploaded as a part of supplementary information.

### CRediT authorship contribution statement

**Subhashree Chatterjee:** Conceptualization, Data curation, Validation, Formal analysis, Visualization, Writing-original draft; **Ritika Jaiswal**: Data curation, *in vivo* experimentation; **Aniruddha Roy**: Validation, Conceptualization, *in vivo* experimentation; **Shibasish Chowdhury:** Docking analysis, Validation, **Sudeshna Mukherjee:** Conceptualization, Funding acquisition, Resources, Supervision. **Rajdeep Chowdhury:** Conceptualization, Supervision, Funding acquisition, Resources, Writing-review and editing.

## Acknowledgement

The authors express their gratitude towards Birla Institute of Technology and Sciences (BITS) Pilani, Pilani campus for providing infrastructural facilities. BITS Pilani for providing student fellowship.

---

**Supplementary Fig. 1. CDDP induced chromatin repression:** (**A-B**) Immunoblots showing expression of H3K27me3 and H3K9me3 in HT29 and HCT116 cells. (**C-D**) Immunofluorescence images of H3K27me3 (red) and H3K9me3 (red) in HT-29 and HCT116. Scale bar 10 μm. **(E-H)** Immunoblots showing expression of EZH2, SUZ12, and JARID2 in HT-29 and HCT116 cells subsequent to CDDP treatment. The data was considered significant (*) if the p-value was ≤0.05.

**Supplementary Fig. 2. Effect of ROS inhibition on cell survival:** (**A-B)** Intracellular ROS levels (DCFDA staining) in HCT116 cells after treatment with CDDP, with or without NAC as measured through flow cytometry (A) and fluorescence microscopy (B). Scale bar 50μm. (**C)** Bar graph showing the intracellular ROS levels (DCFDA staining) after treatment with CDDP, with or without NAC in HT-29 cells as measured through fluorometric analysis. Cell viability as assessed by MTT (**D**) and crystal violet assay (**E**) after CDDP treatment with or without NAC. (**F**) Immunoblot showing expression of JARID2 after CDDP treatment with or without NAC. The data was considered significant (*) if the p-value was ≤0.05.

**Supplementary Fig. 3. Cellular localization of p-P65 post drug treatment: (A-B)** Immunofluorescence images of p-P65 (red) showing the cellular localization of p-P65 under different treatment conditions as mentioned in the figure. (**C-D**) Expression of SUZ12 after siRNA mediated P65 ablation at the mRNA and protein level. (**E**) Protein expression of JARID2 post P65 inhibition**. (F)** Co-immunoprecipitation assay showing the interaction between P65 and SUZ12 in the representative immunoblot. The data was considered significant (*) if the p-value was ≤0.05.

**Supplementary Fig. 4. KDM6A expression in HCT116 cells.** (**A**) The bar graph showing mRNA expression of KDM6A after CDDP exposure with or without NAC. (**B**) Phase contrast images showing a comparison between the cells treated with NAC or ascorbic acid in presence or absence of CDDP. **(C)** Immunoblots showing KDM6A expression after ascorbic acid treatment. (**D**) Immunoblots representing the interaction of KDM6A and PCNA as observed through co-immunoprecipitation assay. The data was considered significant (*) if the p-value was ≤0.05.

**Supplementary Fig. 5. Effect of KDM6A inhibition on cell death and chromatin repression:** (**A-B**) Immunoblots showing expression of KDM6A post inhibition with GSKJ4 or siKDM6A in presence of NAC. (**C-D**) Immunoblots depicting expression of H3K27me3 after GSKJ4 or siKDM6A treatment in presence of NAC. (**E**) Bar graph representing percent cell death as analyzed through flow cytometry using Annexin V-PI staining. (**F**) A pictorial representation of the drug tolerant model development. Cells were divided into 4 groups-untreated control, CDDP exposed (30 μM; 3 days), CDDP rescue (cells were incubated in minus CDDP media for 3 days) and CDDP plus GSKJ4 (GSKJ4 was added after 3 days of CDDP treatment). (**G-H**) Immunoblots showing changes in CD44 and CD133 protein levels following CDDP treatment, with or without GSKJ4, in the presence of NAC, as compared to control. **(I)** Immunoblot showing the expression of CD44 in the *in vivo* tumor samples. The data was considered significant (*) if the p-value was ≤0.05.

**Supplementary Fig. 6. Effect of KDM6A inhibition on normal cells: (A)** Immunoblots showing expression of KDM6A, PCNA after GSKJ4 treatment in HEK293 cells. **(B-C)** Immunoblots showing expression of EZH2 and H3K27me3 after GSKJ4 treatment in HEK293 cells. **(D)** Phase contrast images post GSKJ4 exposure in HEK293 cells.

