## Supplementary Figures for "A switch from histone methyltransferase-EZH2 to demethylase KDM6A activity marks reinitiation of proliferation of drug treated colorectal cancer cells"

**Supplementary Fig. 1.**

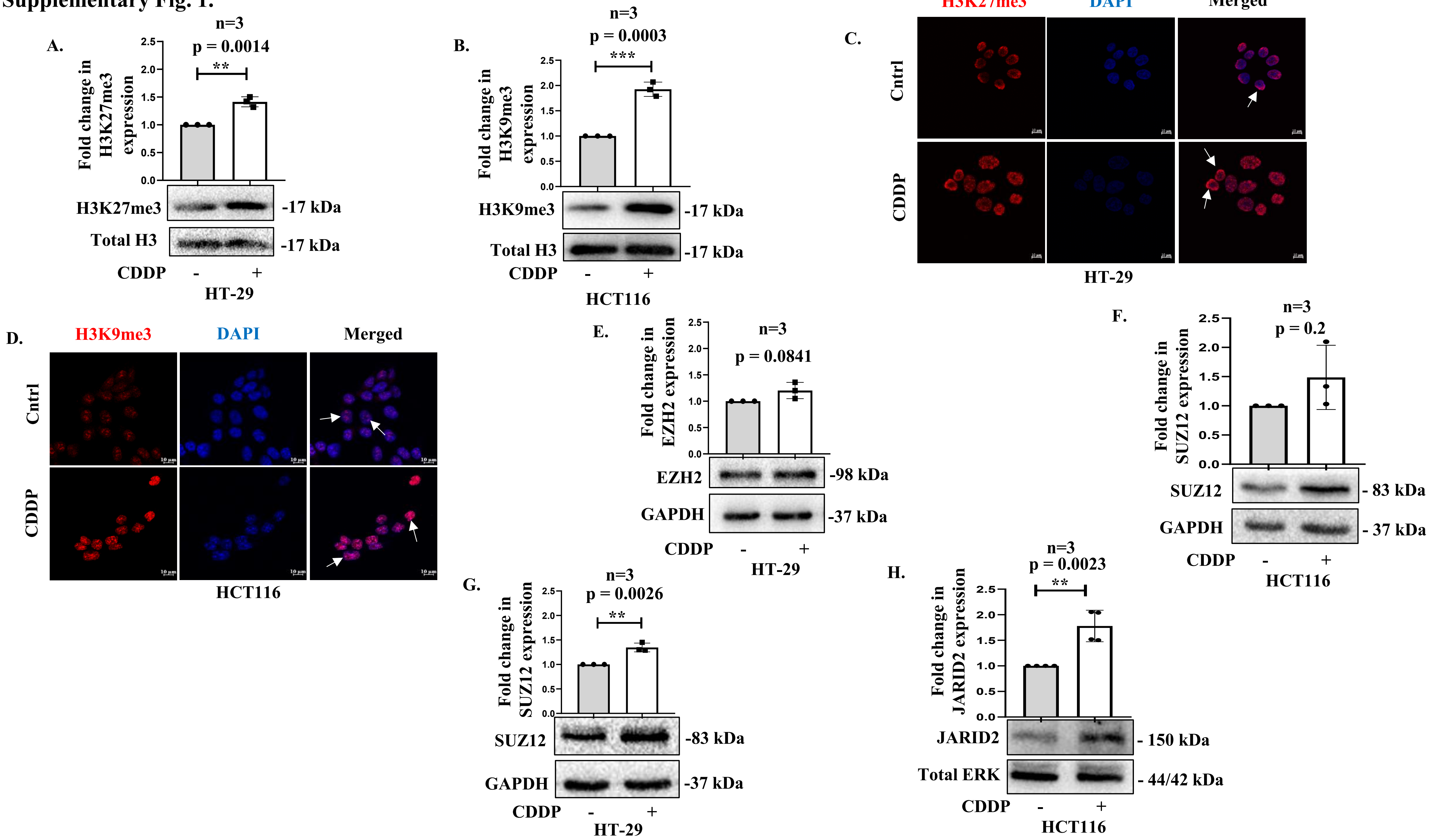

Supplementary Fig. 2.

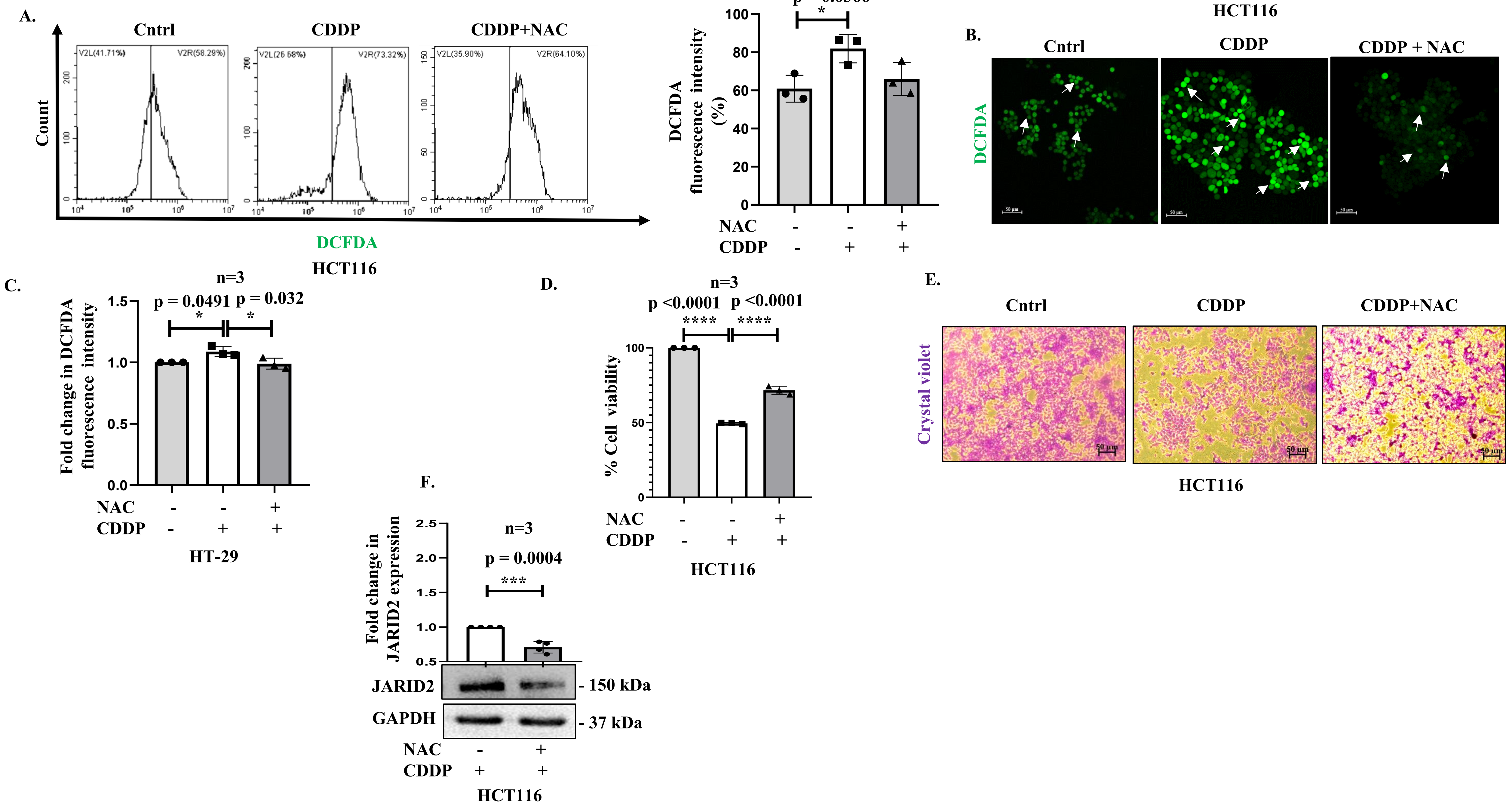

Supplementary Fig. 3.

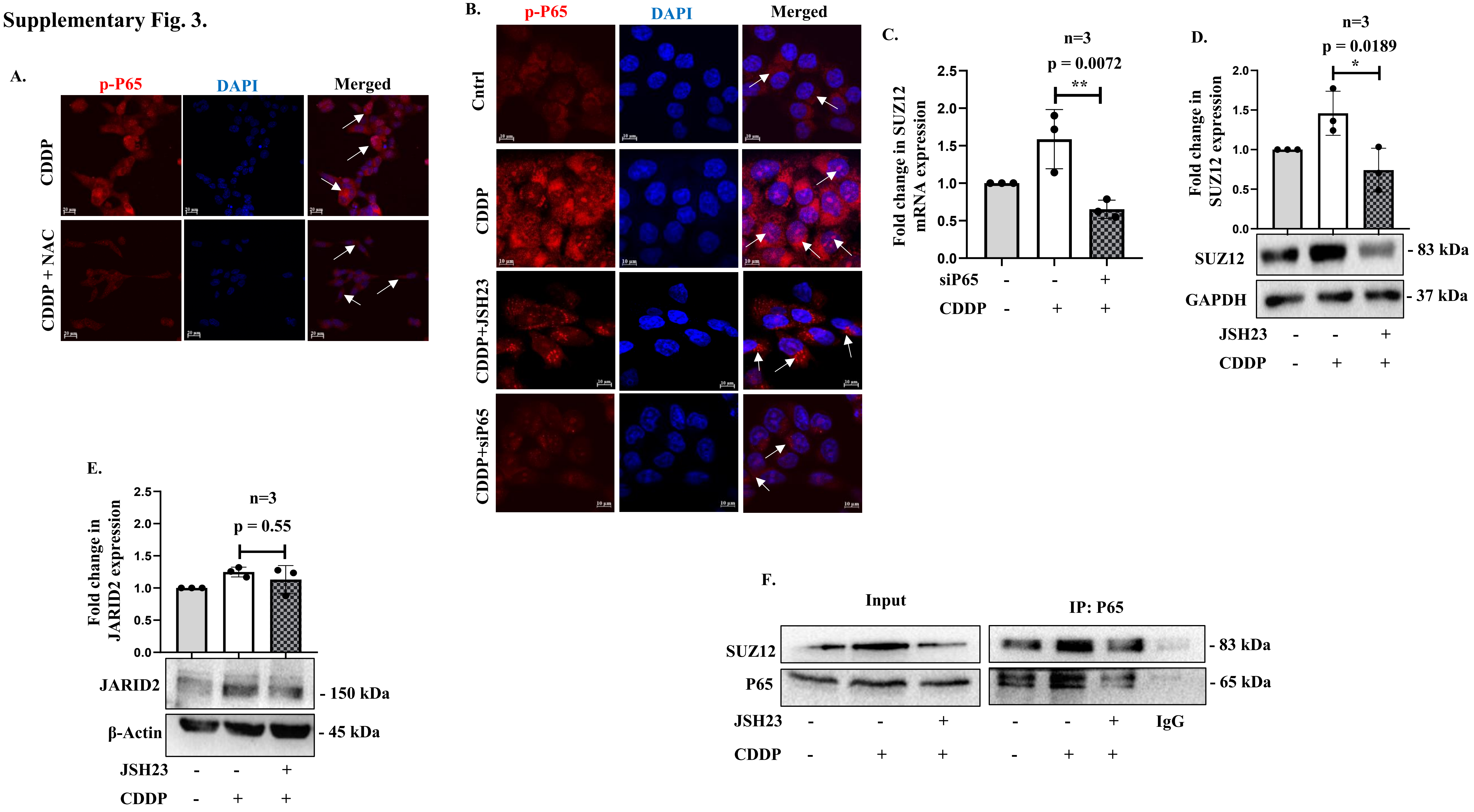

### Supplementary Fig. 4.

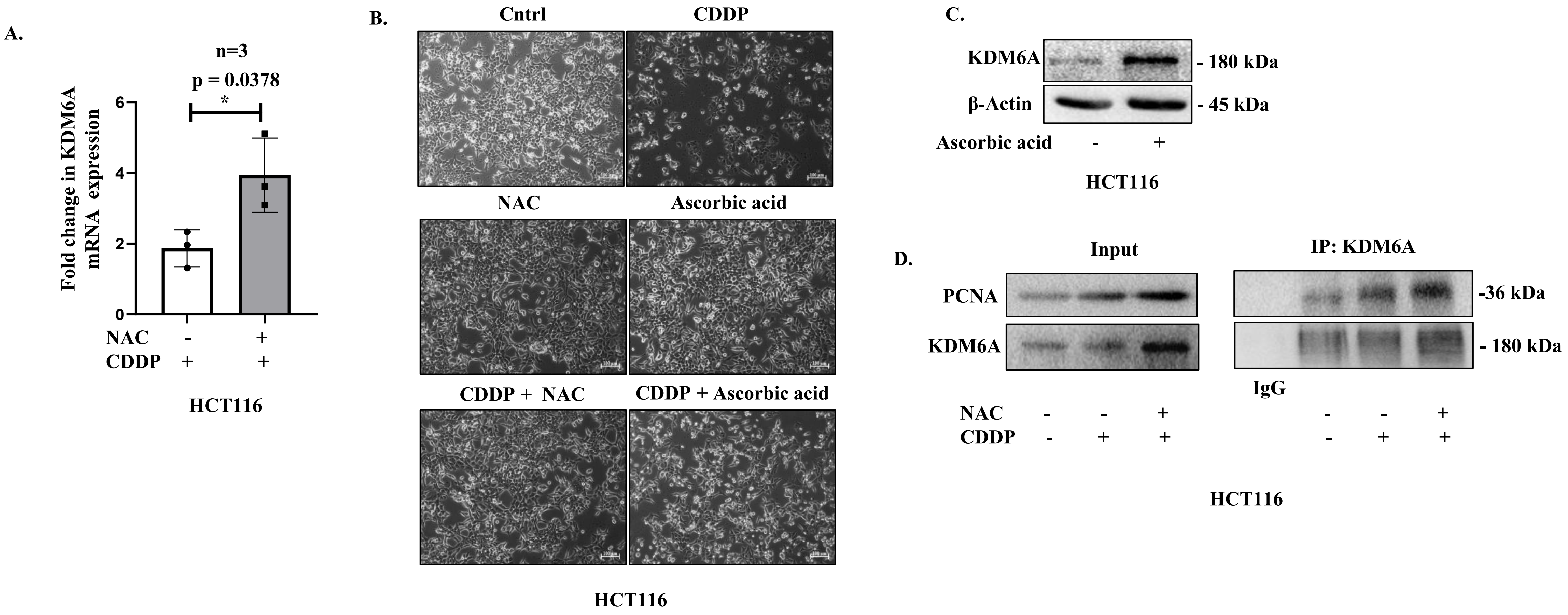

**Supplementary Fig. 5.**

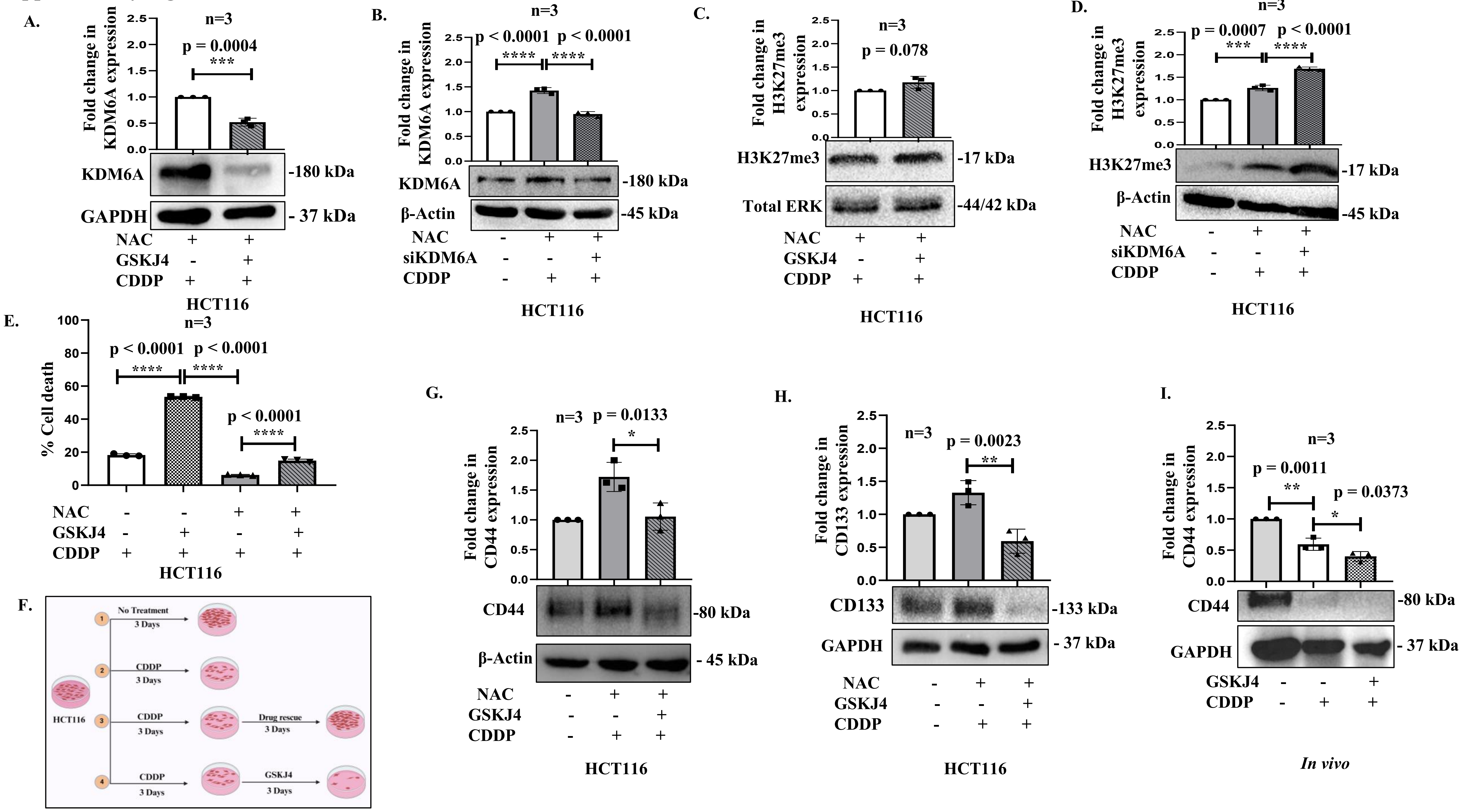

**Supplementary Fig. 6.**

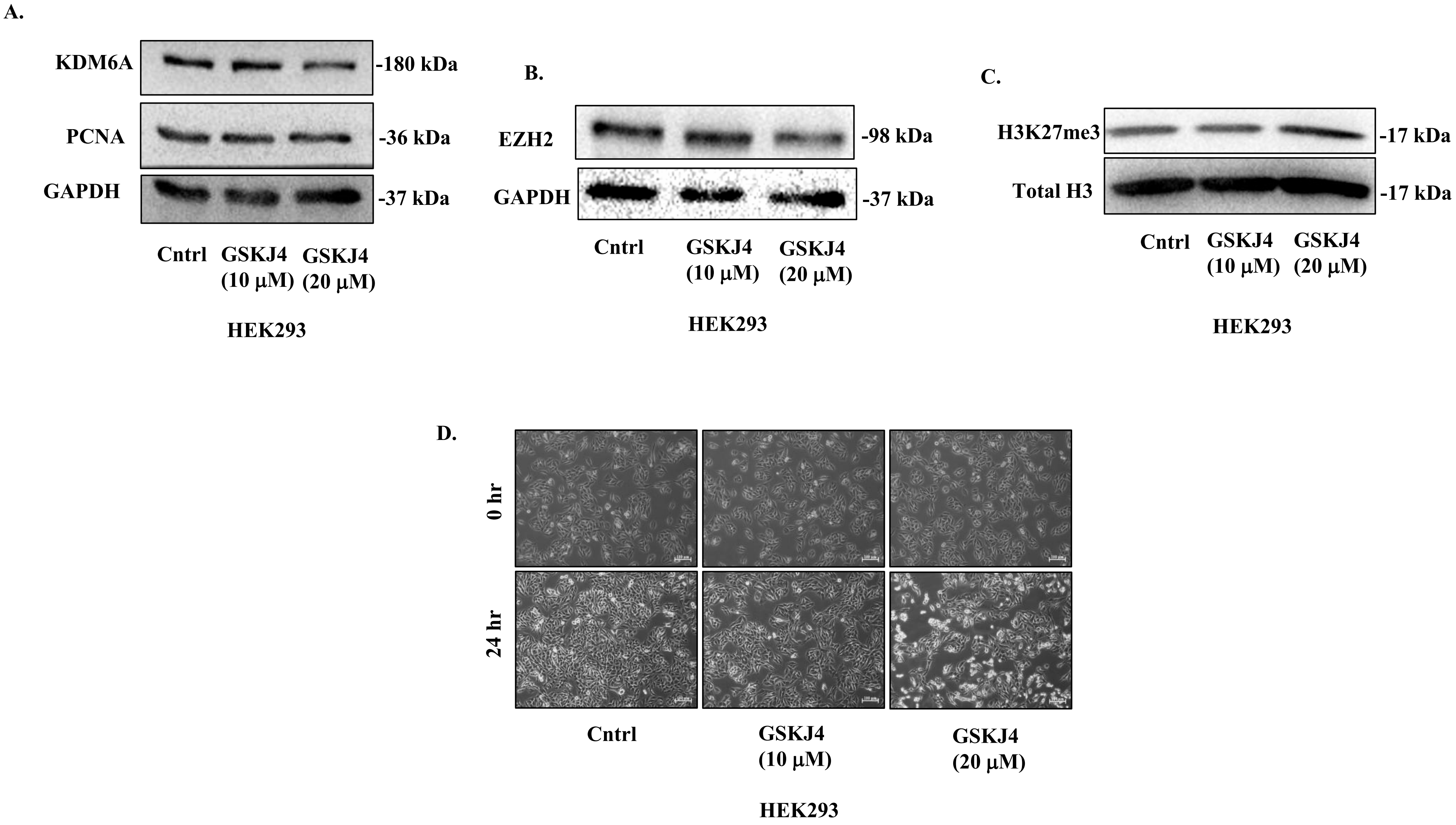
